# Ribo-Pop: Simple, cost-effective, and widely applicable ribosomal RNA depletion

**DOI:** 10.1101/2020.05.19.102293

**Authors:** Mary Kay Thompson, Maria Kiourlappou, Ilan Davis

## Abstract

The measurement of RNA abundance derived from massively parallel sequencing experiments is an essential technique. Methods that reduce ribosomal RNA levels are usually required prior to sequencing library construction because ribosomal RNA typically comprises the vast majority of a total RNA sample. For some experiments, ribosomal RNA depletion is favored over poly(A) selection because it offers a more inclusive representation of the transcriptome. However, methods to deplete ribosomal RNA are generally proprietary, complex, inefficient, applicable to only specific species, or compatible with only a narrow range of RNA input levels. Here, we describe Ribo-Pop (ribosomal RNA depletion for popular use), a simple workflow and antisense oligo design strategy that we demonstrate works over a wide input range and can be easily adapted to any organism with a sequenced genome. We provide a computational pipeline for probe selection, a streamlined 20-minute protocol, and ready-to-use oligo sequences for several organisms. We anticipate that our simple and generalizable “open source” design strategy would enable virtually any lab to pursue full transcriptome sequencing in their organism of interest with minimal time and resource investment.

## INTRODUCTION

Researchers across the biological sciences use RNA sequencing to generate novel insights or test hypotheses. However, in most organisms, ribosomal RNA comprises the majority of the RNA sample, making it economically impractical to analyze sequences of interest such as protein-coding mRNAs and other non-ribosomal transcripts at substantial coverage. Therefore, efficient sequencing requires either capture of the RNAs of interest or depletion of ribosomal RNA.

The most widely used capture method is poly(A) selection, in which the polyadenylated RNAs are purified by hybridization to oligo(dT). Ribosomal RNA, which is not polyadenylated, is greatly reduced in poly(A)-selected sequencing libraries. However, in recent years, interest in the non-polyadenylated transcriptome has grown considerably. Studies which seek to characterize pre-adenylation steps in RNA processing such as nascent transcription and splicing cannot employ poly(A) selecton^1,2^. Furthermore, some transcription events produce RNAs without poly(A) tails. Many non-canonical transcripts, including some classes of enhancer RNAs and anti-sense transcripts, are thought to be unadenylated^3,4^. Moreover, many gene products, such as certain long non-coding RNAs and histone mRNAs, are not polyadenylated, and poly(A) selection does not effectively recover these species^5,6^. Poly(A) selection can also distort abundance measurements of polyadenylated RNAs. RNAs with short poly(A) tails are not captured efficiently by poly(A) selection, and thus use of poly(A) selection can vastly distort gene expression measurements in many contexts^7^. In certain exacting experiments such as measurement of translation or RNA turnover, measurement of polyadenylated RNA rather than the body of the RNA can lead to qualitatively different conclusions^7,8^.

To address these shortcomings, removal of ribosomal RNA, rather than selection of a poly(A) tail, has been introduced in many experimental workflows. Unfortunately, removal of ribosomal RNA is more complicated than poly(A) selection because there is no universal reagent equivalent to oligo(dT) available. Although many protocols and commercial products have been developed to deplete ribosomal RNA or its cDNA derivatives, most of these solutions suffer from various pitfalls that prevent their widespread use. First, each reagent is only applicable to a specific organism or family because hybridization approaches require a high degree of sequence complementarity. For example, as of writing, several commercial kits that deplete rRNA from mammalian samples are available, but none are available for *Drosophila*. Because of the proprietary nature of these kits, researchers working in other organisms cannot benefit from any knowledge used to produce these probe sets in order to design probe sets for their organism of interest. Secondly, even commercial kits are unsuitable for certain types of experiments in their target organism(s). Some of them have narrow RNA input ranges or are not made available as a separate component outside of a sequencing library construction kit, making them difficult to use for novel or exploratory experiments that may deviate from a traditional workflow. Finally, the cost of commercial rRNA depletion reagents is prohibitive for some research groups and prevents their wider adoption.

In summary, we argue that the current lack of versatile and cost-effective rRNA subtraction reagents severely restricts exploration in many areas of RNA biology. Here, we address this problem, by developing an “open source” rRNA depletion solution that is simple to use, cost-effective, and can be easily adapted to any organism of interest. We validate the use of our rRNA subtraction method using RNA purified from *Drosophila* and provide an automated probe design pipeline and pre-designed probes for several organisms. We name our method Ribo-Pop (<u>ribo</u>somal RNA depletion for <u>pop</u>ular use) because it democratizes rRNA depletion, enabling researchers working with any organism or workflow to deplete ribosomal RNA in an efficient and cost-effective manner.

## RESULTS

In an effort to develop high-efficiency ribosomal RNA removal, we decided to test short, biotinylated oligonucleotides for their ability to deplete ribosomal RNA by hybridization and subsequent removal of the rRNA/probe hybrids via pulldown with streptavidin coupled beads. First, we tested how much rRNA depletion is achieved from a single probe/target interaction, in order to determine the minimum number of high affinity probes required to purify unfragmented rRNA. We designed and biotinylated 20 short ∼30mer probes (26 – 32 nt) targeting the small 18S rRNA transcript using parameters developed for microarray probe design and employed in smFISH experiments (Fig. S1)^9,10^. Surprisingly, all probes were able to deplete rRNA to some degree on their own, compared to a control sample containing no probe that was processed in parallel. Each probe was tested individually for its ability to deplete the small 18S rRNA transcript from *Drosophila* total RNA at an estimated 5-fold molar excess in a single-probe depletion assay (Fig. 1A). In the single-probe depletion assay, probe and total RNA were mixed in a standard hybridization buffer (2X SSC, 0.01% Tween-20) and denatured at 70 °C for 5 min. After a brief annealing period at room temperature, the biotinylated probes were captured with streptavidin beads and the remaining rRNA was measured from the supernatant. The average depletion was 3-fold, corresponding to 33% of 18S remaining compared to the no probe control sample (Fig. 1B, Table S1). These results show that many different probes are effective, despite the highly structured nature of the ribosomal RNA molecule.

**FIGURE 1.**
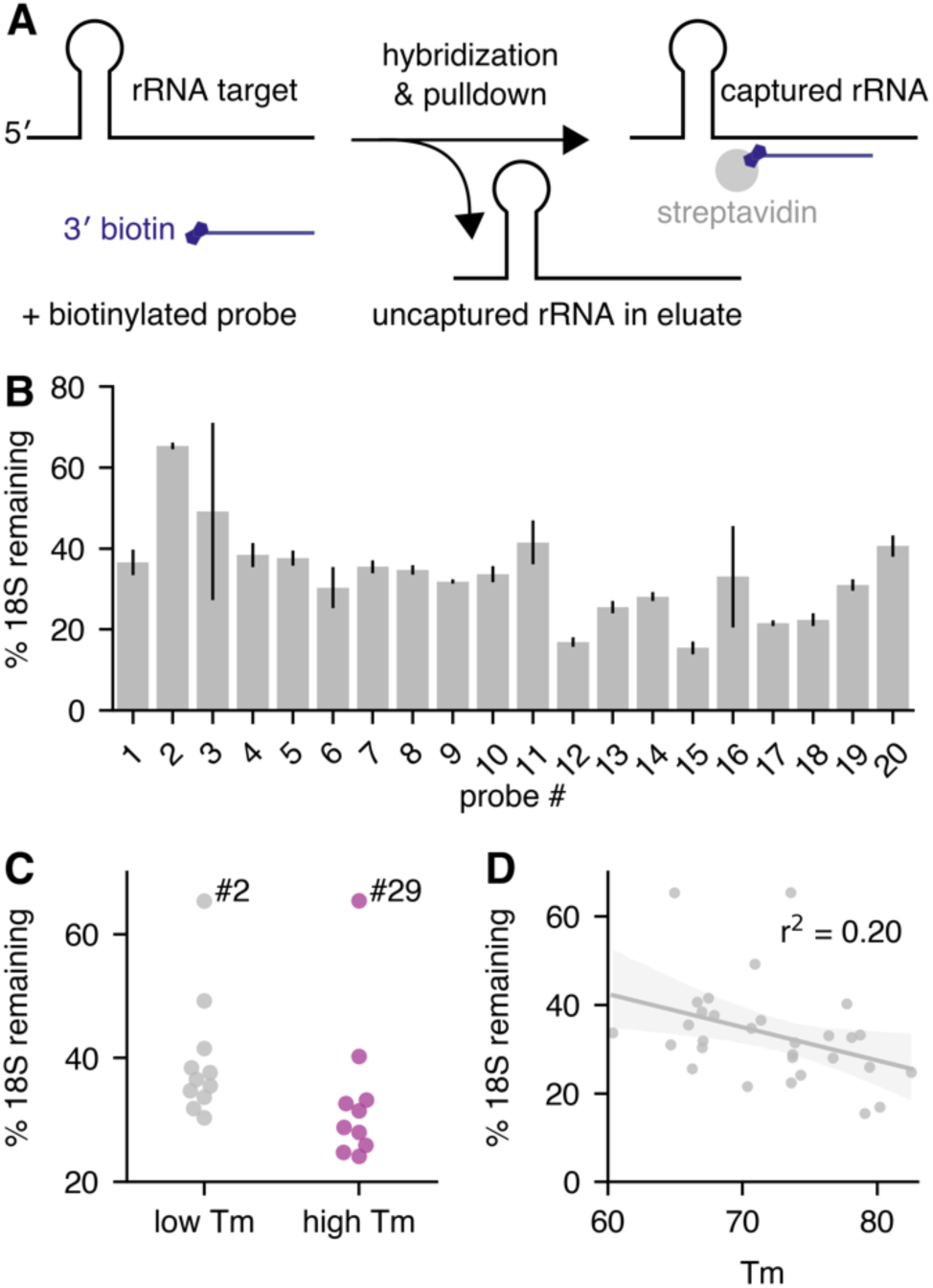
Short anti-sense oligos effectively deplete ribosomal RNA. Oligonucleotides targeting the *Drosophila* small rRNA (18S) were individually tested for their ability to deplete the *18S* transcript from larval total RNA. The percent of remaining 18S was quantified by qPCR. Values are derived from *18S* normalized to *Act5c*, in turn normalized to a non-depleted sample (no probe control). Values are the averages of three replicates of the depletion experiment, each using a different sample of larval RNA. A) Outline of the single-probe depletion assay. A 3′ biotinylated probe targeting a specific site in the 18S is added to total RNA and subjected to hybridization. The target is captured with streptavidin beads and the remaining target is measured from the supernatant. B) The percent of 18S rRNA remaining for all tested oligos of size 26 – 32 nt, arranged 5′ to 3′ by target site. Error bars are standard deviation between separate hybridization experiments, each performed with a different RNA sample. C) Performance comparison between the initial set of low Tm probes (probes #1 – #11, Tm < 71 °C) targeting the left side of the 18S transcript and the second set of high Tm probes (probes #21 – 30, Tm > 71 °C). Two-sided t-test p = 0.12. Probe #2 and probe #29 are outliers, possibly for structural reasons (see figure 2). D) Correlation between the predicted Tm of the probe/target hybrid and the percent of remaining target for the 30 ∼30mer probes tested in the single-probe depletion assay (p = 0.01, Spearman’s correlation).

We chose to employ short ∼30mer probes for several practical and theoretical reasons. Short probes are inexpensive to manufacture at high purity, and they have lower potential for hairpin formation, unwanted dimerization, and off-target binding^11^. However, some studies of microarray probe design suggest that longer probes perform better in that application^11^. We therefore tested whether longer probes designed using our sequence composition criteria would be more efficient at removing rRNA. We found little evidence that longer probes of 46 – 52 nt removed more rRNA than the short probes (Fig. S2A). We hypothesize that the additional nucleotides present in the longer probes make the propensity for self-structure greater and offset the theoretical gain in target affinity. Therefore, we decided to continue using small probes and turned our focus to optimization of probe selection within the 30 nt size range.

We examined our depletion data in order to discover probe properties favorable for ribosomal RNA depletion. Although all probes were able to achieve some degree of depletion, we observed large differences between the best- and worst-performing probes (Fig. 1B). To determine whether thermodynamic properties might explain these differences, we analyzed their relationship with probe efficacy. The correlation between the predicted ΔG for hairpin and homodimer formation of each probe and target depletion were not significant (Fig. S2B – C). In contrast, the predicted Tm of each probe bound to its target was convincingly correlated with target depletion, with higher Tm probes achieving greater depletion on average (r^2^ = 0.17, p = 0.07, Spearman’s correlation) (Fig. S2D).

To test whether probes with Tms that are relatively high (>72 °C) compared to the surrounding sequence area would perform better, we designed probes corresponding to Tm peaks in the first 60% of the 18S target region and tested them for their ability to deplete 18S rRNA. We observed that higher Tm probes did perform better than lower Tm probes on average (3.3-fold vs. 2.7-fold median depletion), although the comparison was not statistically significant (p = 0.12) (Fig. 1C). Nevertheless, when we include all our tested ∼30mer probes in the analysis, Tm and probe performance are significantly correlated (Fig. 1D, r^2^ = 0.20, p = 0.01, Spearman’s correlation). Given this striking correlation, we decided to set high Tm as one of our probe selection criteria.

Given that we planned to select probes with Tm higher than the denaturation temperature (70 °C), we reasoned that annealing would likely begin during the denaturation step, and that the annealing step of the protocol could be eliminated without affecting performance. We tested three different probes with or without the annealing step (Fig. S3). Interestingly, removing the annealing step had no obvious effect on 18S depletion for any of three probes we tested, which ranged in Tm from 65 °C to 80 °C. We therefore eliminated the annealing step from our further experiments and moved directly from the denaturation step to streptavidin binding. Thus, simply by choosing high Tm probes, we were able to create a rapid and robust rRNA depletion strategy.

We then asked whether local or global structure of the target rRNA molecule affects probe performance. We examined the target sites of the 18S probes in the structure of the small ribosomal subunit (Fig. 2)^12,13^. Interestingly, the target sites of successful probes are distributed across the structure with no obvious propensity for single-stranded regions. In fact, many successful probes overlap hairpins that would be expected to require unfolding for the probe to bind. The two worst-performing probes (probe #2 and probe #29, Fig. 1C) overlap with regions of ribosomal RNA expansion segments ES3 and ES6 that are known to interact by base pairing in the 3D structure, forming a helix (Fig. S4)^14,15^. This ES3/ES6 helix comprises 8 consecutive base pairs, but it seems unlikely that this fact alone can explain the poor performance of these probes. Although probe #29 directly interacts with residues involved in the base pairing, probe #2 interacts with residues adjacent to the helix that are not directly involved in base pairing. Furthermore, another probe, #28, targets a hairpin with 8 consecutive base pairs, yet successfully depletes the 18S 4-fold. We postulate that the ES3/E6 long-range tertiary interaction is not completely unfolded after denaturation at 70 °C. In some RNA structures, long-range tertiary interactions form cooperatively at an early stage of the folding pathway and may be more stable than expected^16,17^. If probe #2 and probe #29 are excluded from the high and low Tm probe sets based on their proximity to the ES3/ES6 helix, then difference between high and low Tm performance becomes statistically significant (p = 0.005, Fig. 1C). Taken together, our survey of 18S target sites suggest that successful probes can be chosen by picking probes with predicted Tm above the denaturation temperature and avoiding probes that are likely to contact a high affinity long-range interaction in the rRNA molecule.

**FIGURE 2.**
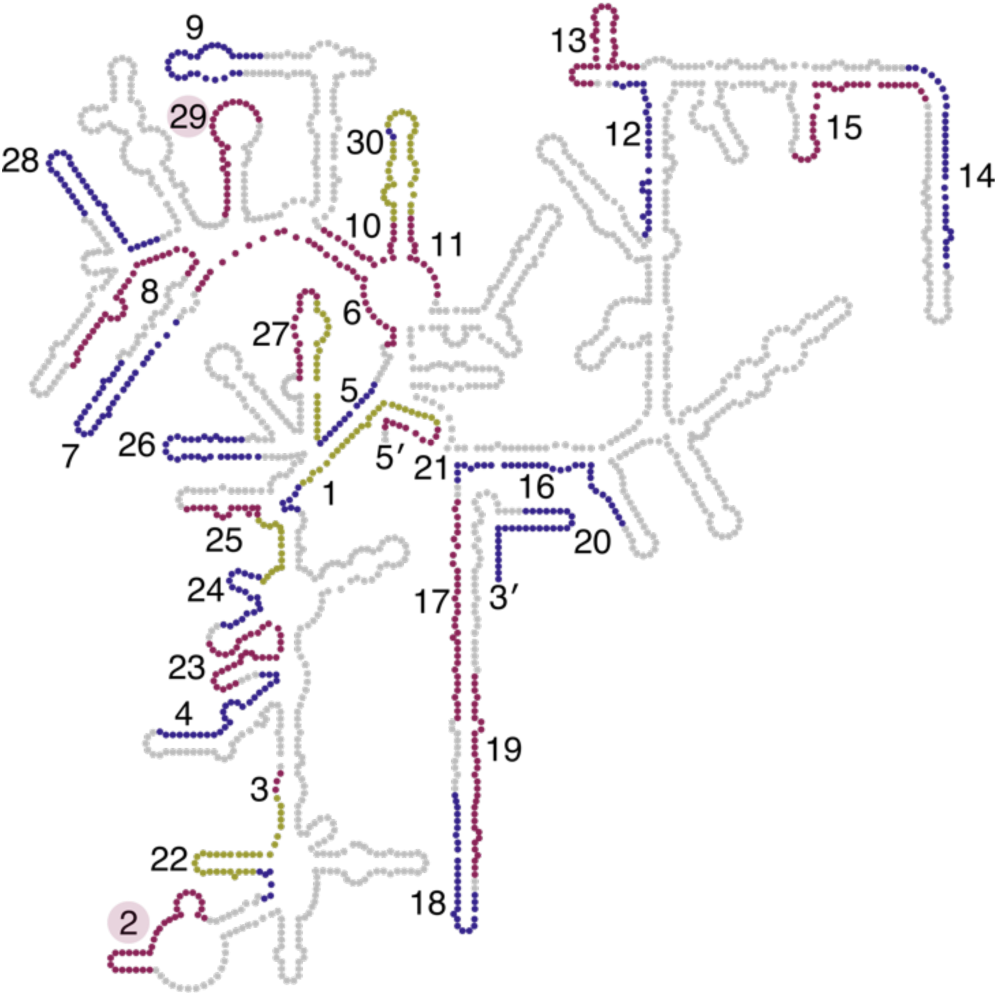
Most 18S structural features are accessible to probe targeting. The target sites of tested probes in the *Drosophila* 18S rRNA is shown. The secondary structure representation is based on data from the 3D structure and constructed using the RiboVision program^12,13^. Some probe target sites are overlapping because we designed a second set of high Tm probes against the 5′ side of the 18S (probes #21 – 30, see figure 1). To distinguish between the individual target sites, we have alternated the colors between red and blue, with the overlapping portions of probe target sites shown in yellow. The labels for the target sites of probe #2 and #29 are highlighted in red.

Using our selection criteria of short, high Tm probes outside of deeply structured regions, we chose a set of 15 probes covering the small and large ribosomal RNA transcripts (5 targeting the 18S, 5 targeting the 28S-L and 5 targeting the 28S-R fragment) spaced evenly along the transcripts to yield a probe density of approximately 1 probe/400 nt. We did not include probes for the smaller transcripts (5S and 5.8S) because we observed that they are not well captured by most RNA-seq protocols and do not contribute substantially to the total rRNA contamination. For the 18S probes, we selected 5 probes from those tested in Figure 1 (probes #12, 18, 21, 24, and 28). For the large subunit 28S fragments, we choose evenly spaced probes conforming to our sequence composition constraints and falling within the top 5% of Tm for potential probes. We also screened probes for potential matches against processed and unprocessed transcripts, using an alignment score cutoff that excludes probes matching more than 15 consecutive base pairs to non-targets^18^. We note that this filter is more stringent than many probe filtering pipelines because we include screening against genomic regions falling within annotated gene boundaries, i.e. intronic sequences found in unprocessed transcripts. Thus, our probe design should allow us to deplete the major ribosomal RNA transcripts with minimal effects on both processed and unprocessed RNA quantification.

We first tested the efficacy and input range of our 15-probe pool using *Drosophila* larva total RNA. We designed both a low-input protocol and a standard protocol to enable the best combination of efficiency and cost-effectiveness for each input amount. Both protocols were able to deplete rRNA efficiently. Using the low-input protocol, we were able to deplete rRNA from 0.5 and 100 ng of total RNA (Fig. 3A). As the typical cell contains 10 – 20 pg of RNA^19^, 0.5 ng represents the amount of RNA that may be obtained from only 25 – 50 cells, which is comparable to a rare cell type experiment.

**FIGURE 3.**
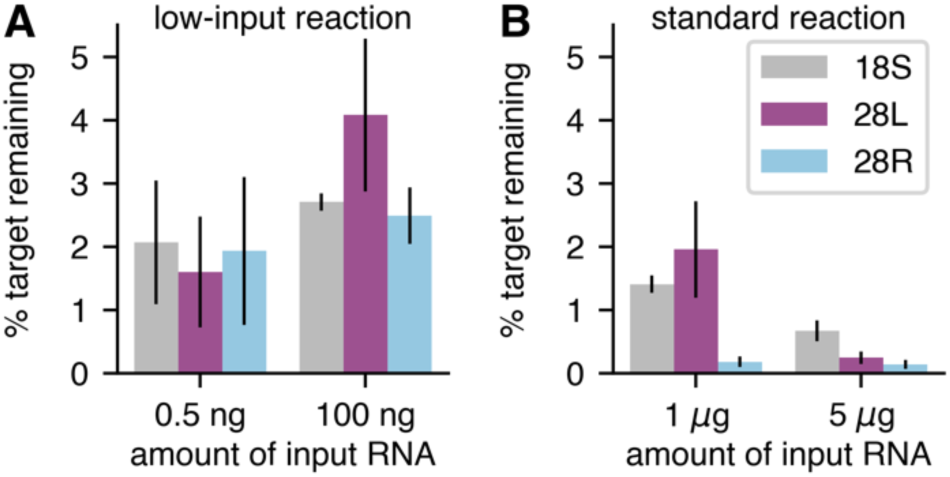
Depletion of rRNA performs well across a range of total RNA input levels. The percent of 18S, 28L, and 28R targets remaining after Ribo-Pop starting from various levels of input RNA is shown. qPCR values for each target were normalized to the *Act5c* level of the same sample and then normalized to an input sample that was not subjected to Ribo-Pop treatment. Samples were subjected to either a low input protocol (0.5 ng or 100 ng of input RNA) or a standard input protocol (1 µg or 5 µg of input RNA). See Methods for a detailed protocol. Error bars are the standard deviation between three independent experiments, each using a different RNA sample. A) Depletion performed with low amounts of total RNA, 0.5 or 100 ng, in a 10 µl hybridization reaction. B) Depletion performed with higher amounts of total RNA, 1 µg or 5 µg, in a 40 or 50 µl hybridization reaction. Sequencing data from the 5 µg samples is presented in Fig. 4.

Using the standard protocol, we were able to deplete rRNA from up to 5 µg of total RNA (Fig. 3B). The difference between the standard protocol and the low-input protocol is simply the amount of probe used and the reaction volume needed to accommodate the additional probe and beads. We also suspected that there would be an effect of target and probe concentration because hybridization is concentration dependent. Indeed, if 100 ng of total RNA is subjected to the standard protocol rather than the low-input protocol, the result is 3- to 9-fold more rRNA remaining after depletion, depending on the target (data not shown). Therefore, we have shown Ribo-Pop is effective and applicable over at least a 10,000-fold range if the correct protocol is selected.

It was important to know what improvement in effective sequencing depth could be achieved by applying Ribo-Pop to a typical RNA sequencing experiment. We subjected 5 µg of *Drosophila* total larval RNA to Ribo-Pop depletion and prepared sequencing libraries from the remaining RNA. Non-depleted RNA-seq libraries (“input”) were prepared in parallel. After sequencing, we mapped the reads to all annotated transcripts and counted the sum of reads counted per transcript class (Fig. 4A). Ribo-Pop depletion decreased the percentage of reads assigned to rRNA transcripts from an average of 97% of reads in the input libraries to an average of 27% of reads in the depleted libraries. Concurrently, depletion greatly increased coverage on non-rRNA transcripts. Protein-coding transcripts increased from 2.7% to 61.9% of reads and ncRNAs increased from 0.4% to 7.1% of reads. We also added synthetic spike-in RNAs to the input samples before depletion, and coverage on these spike-ins increased from 0.1% to 2.8% after depletion. Hence, Ribo-Pop depletion increased the effective depth of the sequencing experiment more than 20-fold.

**FIGURE 4.**
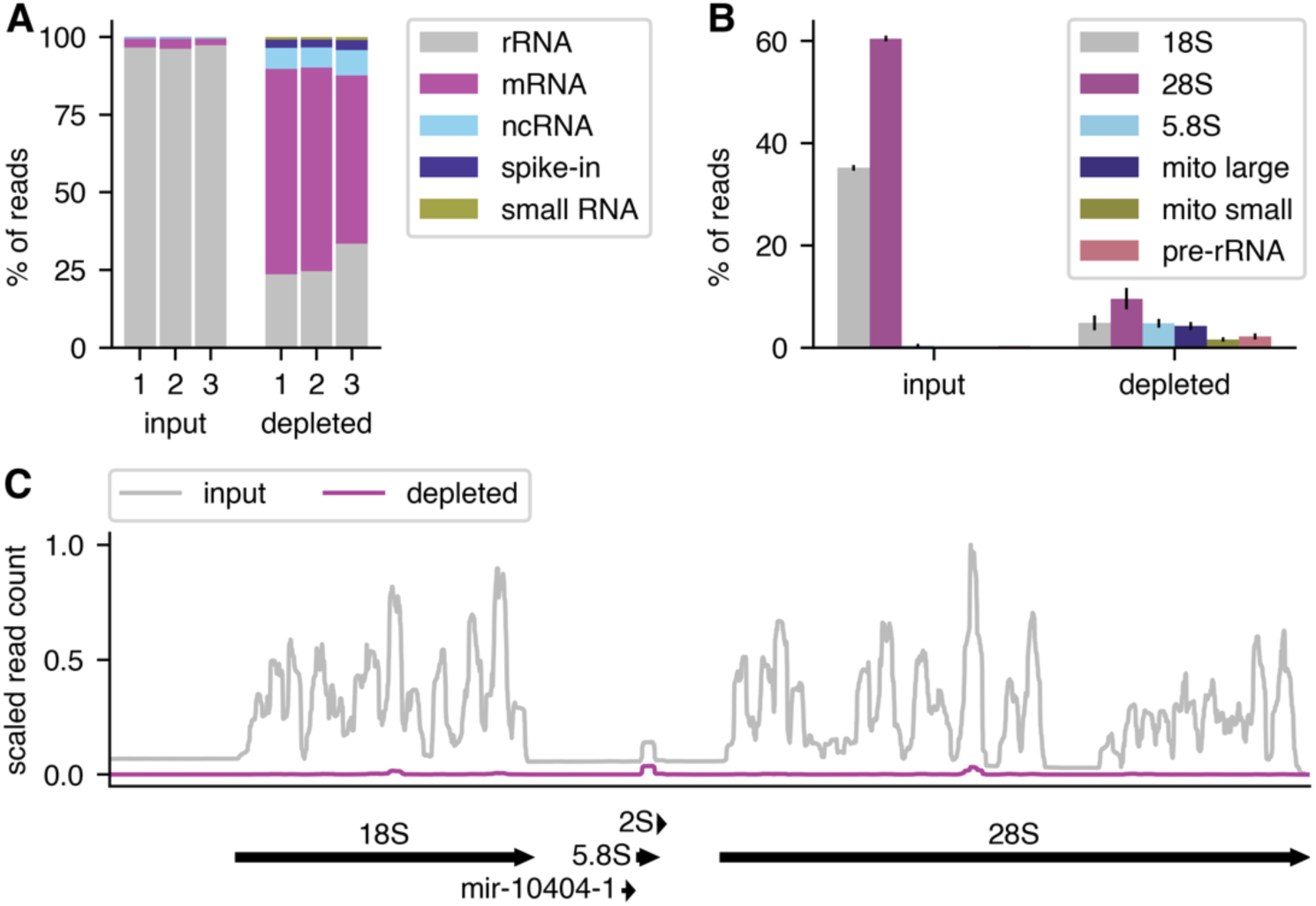
Ribo-Pop drastically increases effective sequencing depth. The effectiveness of Ribo-Pop depletion was assessed by RNA sequencing of input and depleted libraries prepared from *Drosophila* larval RNA. A) The percent of reads assigned to each transcript class (biotype), is plotted for each of the three replicates of the input and subtracted RNA samples. B) The percent of reads assigned to each type of ribosomal RNA is shown for the input and depleted samples. Error bars represent standard deviation. C) RNA-seq coverage of a representative pre-rRNA locus (FBgn0267507) and its cleavage products in input and depleted samples. Reads in each library were scaled by the synthetic spike-in counts to allow direct comparisons between libraries. The average of three replicates for each sample type is plotted.

We then examined the remaining rRNA contamination in our depleted libraries. The remaining 27% of rRNA contamination is comprised of 9.6% 28S, 4.8% 18S, 4.2% mitochondrial large rRNA, 4.8% 5.8S rRNA, 2.2% pre-rRNA, 1.6% mitochondrial small rRNA, and 0.04% 5S (Fig. 4B). Thus Ribo-Pop depleted the targeted 18S and 28S RNAs down to levels similar to the minor rRNA contaminants (mitochondrial rRNAs, pre-rRNA, 5S and 5.8S), which together make up around 1% of reads in the input libraries. RNA depletion occurs relatively evenly across the targeted rRNA transcripts (Fig. 4C), suggesting that the remaining contamination is widely distributed. It is important to note that the observed increase in the percent of reads assigned to non-targeted RNAs can be explained by the increased sequence space available after depletion of the 18S and 28S transcripts. Likewise, examining the percent of target RNAs remaining is not an accurate measure of their fold change upon depletion. The actual fold change can be estimated by normalizing read counts to synthetic spike-in RNAs that were added before the depletion step. Using this normalization method, we find that the 18S and 28S decrease to approximately 0.5% and 0.6% of their input RNA levels, respectively, in agreement with our qPCR results. Hence, Ribo-Pop greatly depletes its targeted rRNA transcripts.

To test the specificity of our probe pool, we compared poly(A)-primed sequencing from depleted samples to poly(A)-primed sequencing of input samples. We chose to use poly(A)-primed data for this analysis because the low depth of sequencing in non-poly(A) primed, non-depleted input samples precludes quantification of most genes. We found that Ribo-Pop depletion had little effect on our gene quantification. The correlation between input and depleted RNA levels for mRNAs and ncRNAs was high (Pearson r^2^ = 0.97, Fig. 5A). We performed differential expression analysis and found that at an adjusted p-value cutoff of 0.01, 30 non-rRNA genes were called as upregulated by subtraction and 217 non-rRNA genes were downregulated, with a median fold-change of 1.9- and 2.2-fold, respectively (out of a total of 14,852 quantified genes, Fig. 5B). The downregulated genes do not have more sequence complementarity to the probes than all other genes, suggesting that our probe specificity filtering approach was successful (Fig. 5C). Therefore, depletion does not substantially bias gene quantification.

**FIGURE 5.**
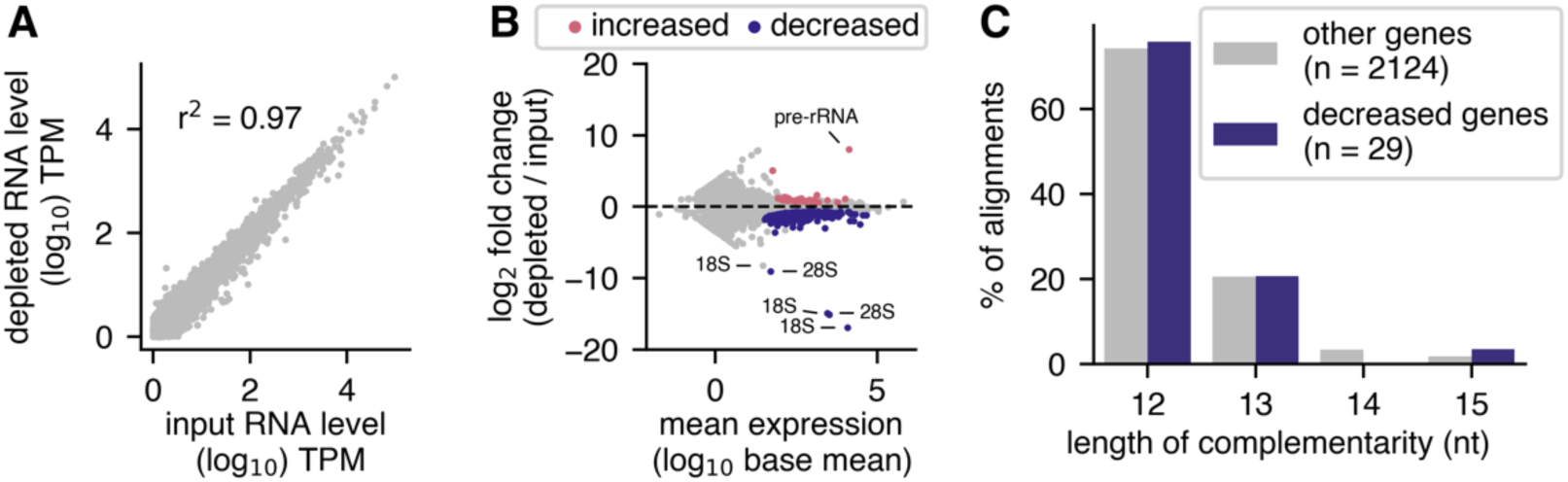
Ribo-Pop has minimal effects on expression measurements of non-rRNA genes. RNA-seq libraries prepared after Ribo-Pop depletion were compared to libraries made from non-depleted input RNA. To allow sufficient coverage for analysis, libraries were prepared using a poly(A)-primed method (QuantSeq). A) Scatterplot and the Pearson’s correlation between input and depleted RNA levels for mRNAs and ncRNAs. The RNA levels are the averages of three replicates for each condition, filtered to include only genes with an average of at least 1 transcript per million (TPM). B) MA plot for the comparison of Ribo-Pop depleted vs. input libraries analyzed with DESeq2^20^. The data was obtained from three independent RNA samples in which each sample was used to construct one input library and one depleted library. Genes with an adjusted p-value <0.01 for differential expression are highlighted in pink if increased upon depletion or blue if decreased upon depletion. Some notable rRNA genes are indicated. C) Stretches of complementarity between the Ribo-Pop probes and the transcriptome were identified by BLAST^21^ and classified as overlapping a gene decreased upon Ribo-Pop depletion (decreased) or overlapping a gene not decreased upon depletion (other). Only alignments with 100% identity are displayed.

Finally, our goal in developing Ribo-Pop was not merely to create an rRNA removal protocol for ourselves, but to provide a general solution for any researcher to apply to their organism(s) of interest. To this end we wrote a fully-automated pipeline for probe selection, which includes selection of evenly-spaced, high Tm probes and specificity checking (Fig. 6A). The pipeline is implemented in Snakemake^22^, which makes the entire probe design process deployable with a single command. We also pre-designed probe sets for common model organisms, where possible generating a single probe set to target multiple organisms (Table S2). For example, the pipeline designed a probe set targeting both *S. cerevisiae* and *S. pombe* yeast rRNA by finding regions of identity between each target and then choosing probes corresponding to our selection criteria (Fig. 6B). In conclusion, we anticipate that Ribo-Pop will prove a versatile and cost-effective tool to enable full transcriptome sequencing for researchers working in many different systems.

**FIGURE 6.**
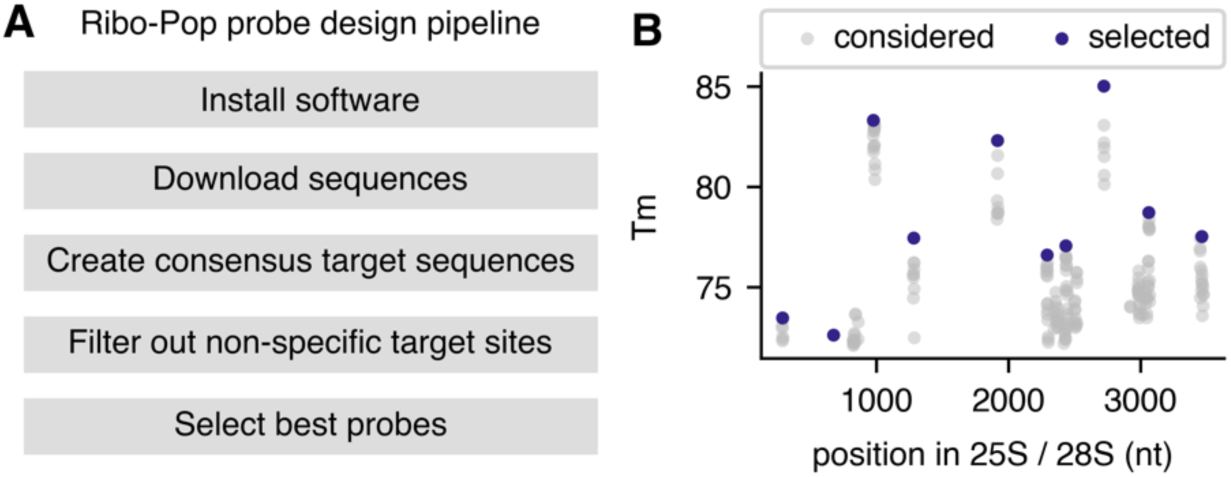
The Ribo-Pop probe design pipeline can choose probes for other organisms. A) The Ribo-Pop design pipeline. After software installation, the remaining steps can be automatically deployed. B) An example output figure of the Ribo-Pop design pipeline after running the pipeline on the major rRNA transcripts of *S. cerevisiae* and *S. pombe* yeasts. The probe selection process for the large rRNA transcript is shown. Grey dots indicate possible probe placement sites based on conservation and sequence composition. Blue dots indicate the selected probes sites based on a desired probe set size of 10.

## DISCUSSION

We have described a simple, effective, and affordable strategy for ribosomal RNA removal that can be extended to any organism with a sequenced genome. Using only 15 short ∼30mer probes, we were able to reduce rRNA levels greatly and increase the percent of non-rRNA reads from 3% to 73%. Furthermore, by testing individual probes, we elicited information about the importance (or lack thereof) of probe length, Tm, and structural effects. We argue that these insights and the computational design pipeline that we have developed will be helpful to any researchers wishing to design their own probes and may even have relevance beyond rRNA removal for other types of RNA pulldown experiments, such as viral RNA purification.

Ribo-Pop has low start-up costs compared to many other non-commerical rRNA subtraction methods. By testing individual probes, we were able to identify features of successful probes and assess how many probes would be needed to obtain the desired results, without resorting to an unnecessarily costly excess of probes and streptavidin matrix. The oligos can be ordered biotinylated for around $1000 - $2000, but thereafter used at a relatively low cost per reaction. As described here, oligos can also be ordered unmodified and biotinylated in-house for a start-up cost of around $50 per probe set and the additional cost of the TdT enzyme and ddUTP for a total start-up cost of around $150. The low costs and ease of adding or subtracting probes from the mix will make it simple for researchers to test and optimize custom mixes if needed.

A researcher wishing to deplete as much rRNA as possible may wish to add more probes to target the other contaminants or to tile across the 18S and 28S more densely. However, one should consider whether such a strategy is optimal from a cost-benefit perspective. To do so would more than double the size of the probe set and increase the cost of the experiment more than 2-fold. If the expanded probe set were to decrease rRNA-mapping reads to 5%, then the space available for non-rRNA reads would increase from 73% to 95%, an increase of only 30%. In other words, greatly increasing the target space of the probe set and the cost per experiment is expected to increase the effective sequencing depth only modestly. For this reason, we chose to remain focused on the major contaminants, the processed rRNA transcripts of the small and large cytosolic ribosome. However, our probe design pipeline can accept user parameters to design more densely packed probes or to include more targets if desired.

One potential pitfall of our 15 oligo probe set is that it would not be expected to perform as well on fragmented rRNA. Use of a larger probe set entails a higher cost per reaction, but would be predicted to perform better on fragmented samples. Another option for highly fragmented RNA may be an enzymatic depletion method where rRNA is bound by tiling DNA probes and then subjected to RNAse H digestion. The RNAse H treatment is sometimes preceded by a reverse transcriptase step to extend the DNA/rRNA hybrids and is always followed by a DNAse treatment to remove the DNA probes^23,24^. These steps lead to a longer depletion protocol and possibly more opportunities for RNA degradation or non-specific RNA targeting than Ribo-Pop. Furthermore, RNAse H protocols are typically applied to total RNA < 1 µg, presumably due to the cost of reaction scale up for the multiple enzymatic steps. Nevertheless, RNAse H methods may be more suitable than pulldown-based methods when dealing with low-input, highly fragmented RNA samples.

During preparation of this study, a handful of similar protocols using short probes were published using biotinylated oligos of approximately 50, 40, or 30 nt, followed by streptavidin pulldown^25–27^. Two of these studies were validated with relatively small samples (100 ng) and used 2-3 rounds of depletion on the same sample, resulting in a more laborious protocol that uses many more moles of streptavidin per ng of total RNA than Ribo-Pop^25,26^. One of these studies, which used 88 40-nt oligos and 60-fold more streptavidin than Ribo-Pop, achieved higher rRNA depletion levels than Ribo-Pop^26^. We believe that the difference is due to their larger probe set, which includes both denser tiling of the 18S and 28S and inclusion of additional target RNAs. Their expanded probe set and greater decrease in rRNA contamination comes at much higher cost per ng of total RNA that we found unjustified in our experiments. The other study used 90 times more streptavidin despite using only 12 50-nt probes^25^. We do not know the reason they used such high molar excess of probes and streptavidin, but we speculate it may be related to the relatively low salt concentration and the inclusion of formamide in their hybridizations, conditions which caused reduced depletion efficiency in our pilot experiments (data not shown). Therefore, we believe that Ribo-Pop is faster and more economical than these related methods.

The most recently published protocol, the DIY method, which used 21 30-nt probes is most similar to ours in probe length, number, and depletion protocol^27^. In that study, the authors chose to focus on a design that would be able to target eight different bacterial species. Thus, they allowed mismatches between the probe and target and selected probes with a lower Tm than the Ribo-Pop pipeline. They achieved similar depletion levels to Ribo-Pop, although a direct comparison is difficult because bacteria lack 5.8S and mitochondrial rRNAs. Impressively, they were able to deplete rRNA from highly distant species. The Ribo-Pop probe design pipeline does allow the user to design probe sets targeting multiple species. However, it does not currently support probe design at alignment sites containing mismatches between the target species. Given the prevalence of short stretches of nucleotide identity suitable for probe design and the relatively low cost of creating additional Ribo-Pop probe sets, we do not feel that this feature is required at this time. Users that need to deplete rRNA from a pool of distantly related microbial communities may find the DIY design better suited to their needs, although another option could be to use Ribo-Pop to select high Tm probes from subsets of more closely related species and then pool them to create a probe set against a very diverse set of species. It is difficult to know whether inclusion of more high Tm probes or fewer low Tm probes would lead to better rRNA depletion in this type of experiment. In any case, either of these tools should provide an effective open source option to design rRNA removal probes that target multiple species.

Ribosomal RNA removal reagents should be available to the entire scientific community, not only those that happen to work with the organisms best served by the biotechnology industry. The cost of RNA sequencing has been decreasing rapidly over the last ten years^28^. While previously the cost of the sequencing reaction itself was the main financial barrier to performing these experiments, now with falling sequencing costs, sample preparation costs can become a limiting factor^28^. The use of a commercial ribosomal RNA extraction kit can easily more than double the total sample preparation cost. As statistical power and the ability to draw biologically meaningful conclusions grow with the addition of more replicates and more biological conditions^29^, high sample costs relative to sequencing costs discourage discovery-driven science. Furthermore, the lack of readily available solutions for some model organisms (such as *Drosophila*) and other organisms of ecological interest effectively prevent or severely limit the application of many RNA sequencing protocols in these systems. This discrepancy further widens the gap between often better-funded mammalian work and frequently underfunded work on invertebrate models. We argue that rRNA depletion is the most appropriate approach to informative RNA enrichment for many biological questions, and that “open source” reagents as cost effective and readily available as oligo-dT for poly(A) selection should be made available. We believe that the Ribo-Pop method will help bridge this resource gap.

## MATERIALS AND METHODS

### Probe design for the Drosophila rRNA transcripts

We first created an alignment of the *Drosophila* 18S from the three annotated non-pseudogene variants (FBtr0346874, FBtr0346882, FBtr0346878) using MAFFT^30^ with the --auto parameter. A consensus sequence was created in which mismatches and internal gaps in the alignment were replaced by Ns. We input the 18S sequence into the Oligostan.R script (designed for smFISH probe design)^9^ with the default parameters (min length = 26, max length = 32, ΔG_min_ = -36, ΔG_max_ = -28). For the longer 46 – 52mer probes, we changed the desired probe length to these ranges and set the ΔG_min_ = -56 and ΔG_max_ = -48. We then screened the returned probes to remove probes with 5 or more consecutive identical nucleotides, 4 or more consecutive As, or 4 or more consecutive Cs in the first half of the probe sequence, as previously recommended^9,31^. We also removed probes with GC content <40% or >60%^9,31^ and probes that overlap with Ns in our consensus sequence of 18S variants. We chose 20 probes of length 26 – 32 and 5 probes of length 46 – 52 nt from these sets for our initial tests. We later chose 10 high Tm probes corresponding to Tm peaks on the left side of the 18S. The high Tm probes have otherwise the same length and thermodynamic property ranges (predicted ΔG for homodimer and heterodimer formation) and meet the same sequence composition criteria as the first set of 20 short probes. The predicted Tm of each probe/target interaction was calculated using Biopython^32^ with nearest-neighbor values measured for RNA/DNA hybrids^33^, the DNA concentration set to 250 nM, and salt correction applied^34^ for 300 mM Na^+^. ΔG estimates were obtained from Primer3^35^ using a Na^+^ concentration of 300 mM.

After our initial tests we decided to enforce additional probe selection criteria. We limited our selection to probes with a predicted ΔG of hairpin formation > -3 kcal/mol and a predicted ΔG of homodimer formation or heterodimer formation with other probes to > -10 kcal/mol. We also incorporated screening for probes with potential off-target binding using BLAST^21^ alignment. We built a custom BLAST database containing both spliced transcripts and introns with 40 nt of flanking sequence, then aligned the reverse complement of our probe candidates to the transcript database. To ensure we detected short regions of homology, we used the blast command “blastn -task blastn-short -dust no -soft_masking false -evalue 50”. We removed any probes that had an alignment bitscore >32, which catches probes with 16 or more contiguous matches to a non-target transcript or a longer alignment with some mismatches. This filter is similar to one used for microarray probe selection^18,36^.

The five probes targeting the 18S rRNA that were included in the final mix were picked from the initial set of 30 tested probes after checking that they met our these additional thermodynamic and specificity criteria. For probes targeting the *Drosophila* 28S rRNA, we made a consensus sequence of FBtr0346876 and FBtr0346885 and selected probes of 26 – 35 nt that met the same constraints and fell within the top 5% of Tm for available candidates in a 200 nt moving window. We excluded probes that overlap regions involved in long-range base pairing: between ES3 and ES6 in the 18S and between helix 22 and helix 88 in the 28S.

### Probe design for additional species with the Ribo-Pop probe design pipeline

To enable streamlined design of additional probe sets, we wrote a pipeline to automate all tasks of probe design. The pipeline builds a consensus sequence for each target, so that multiple transcript variants and/or targets from multiple organisms can be used as input. The consensus sequences are masked^37^ to prevent selection of probes overlapping repetitive regions. Candidate probes are then created in the user-defined size and Tm range and filtered to remove probes with undesirable sequence composition or thermodynamic properties, as described in the previous section. The pipeline automates building of the BLAST transcript database and screening of probe candidates for off-target binding. Finally, the pipeline selects candidate probes at Tm peaks that are as evenly spaced along each target as possible. The pipeline is available at https://github.com/marykthompson/ribopop_probe_design. The pipeline is implemented in Snakemake^22^.

### Probe biotinylation and purification

Unlabeled oligos were ordered from Sigma Aldrich or Invitrogen with desalting purification and resuspended in probe resuspension buffer (10 mM Tris 8, 0.1 mM EDTA). Probes were biotinylated with terminal deoxynucleotidyl transferase (TdT, ThermoFisher) and dideoxy-UTP-biotin (biotin-11-ddUTP, Jena bioscience) as previously described^10^. To biotinylate single probes for testing, we set up 5 µl reactions containing 100 pmol unlabeled oligos, 500 pmol ddUTP-biotin, and 4U TdT. To biotinylate oligo pools, we set up 15 µl reactions containing 1000 pmol mixed unlabeled oligos, 3000 pmol ddUTP-biotin, and 12U TdT. Reactions were incubated overnight at 37 °C in a PCR machine and terminated by heat inactivation at 70 °C for 10 min. 5 µl reactions were purified with the ZR-96 Oligo Clean & Concentrator and 15 µl reactions were purified with the Oligo Clean & Concentrator kit (Zymo Research) and eluted with probe resuspension buffer. Probe concentration was determined using duplicate measurements from a NanoDrop ND-1000 instrument using the ssDNA setting and molarity was determined using the A260 value and the nearest-neighbor extinction coefficient estimated from the OligoEvaluator tool (Sigma Aldrich). Biotinylation efficiency was assessed by running an aliquot of the biotinylated oligo alongside unbiotinylated oligo of the same sequence and 10 bp DNA ladder (ThermoFisher) on a 20% TBE-Urea PAGE gel (SequaGel, National Diagnostics) and staining with SYBR Gold (ThermoFisher). Images of the gels were captured with a LI-COR Odyssey Fc.

### *Drosophila* culture and RNA extraction

*Drosophila melanogaster* wild type strain Oregon-R flies were raised on standard cornmeal-agar medium at 25°C. Three samples of 40 wandering third instar larvae each were collected and RNA was extracted with Trizol^38^. Prior to rRNA depletion, 40 µg aliquots of larval RNA were treated with 8U of TURBO DNAse (Thermofisher) for 30 min at 37°C, then purified with the RNA Clean & Concentrator kit (Zymo Research) and eluted with water.

### Quantitative PCR

cDNA synthesis was performed with UltraScript Reverse Transcriptase (PCR Biosystems) in 10 µl reactions containing a mix of RNA, random hexamer (5 µM), anchored oligo-dT primer (1 µM, TTTTTTTTTTTTTTTTTTTTVN, Jena Bioscience), buffer, and 100U UltraScript enzyme. Reactions were denatured 5 min at 70°C with RNA and primers alone, then placed on ice for 5 min. The remaining components were then added and the reactions were incubated for 30 min at 42°C, followed by 10 min at 85°C. cDNA was amplified with qPCRBIO SyGreen Mix Lo-Rox (PCR Biosystems) according the manufacturer’s instructions in 10 µl reactions on Bio-Rad CFX96 instrument. The qPCR primer sequences were taken from a previous study^39^ and are listed in Table S3. Standard curves with cDNA dilutions were routinely performed to ensure the amplification efficiency was close to 100% and no RT controls were also assessed to verify the success of DNA removal from the RNA samples. The amplification Ct values were called automatically by Bio-Rad CFX Manager. Fold changes are derived from the ΔΔCt method. Each target Ct value was first normalized by *Act5c* (ΔCt), followed by normalization by a control sample (either no probe or input RNA, as indicated in the appropriate figure legend) to produce ΔΔCt.

### Single-probe depletion assay

Biotinylated probes were brought to 0.2 µM in probe resuspension buffer (10 mM Tris 8, 0.1 mM EDTA) and used to deplete rRNA from 200 ng of larval total RNA. To minimize the effect of pipetting error on probe efficacy measurements, each replicate uses a different sample of larval RNA and an independent dilution of biotinylated probe. Each probe was hybridized in a 10 µl reaction containing 200 ng total RNA and 0.5 pmol of probe in Ribohyb buffer (2X SSC, 0.01% Tween-20). Reactions were heated to 70 °C for 5 min in 1.7 ml tubes in a thermomixer and then placed at room temperature for 10 min before capture with Dynabeads MyOne Streptavidin C1 (ThermoFisher). To prepare beads for capture, 2 µl beads per sample were washed 3X in MyOne B & W buffer + 0.01% Tween, 2X in Dynabeads buffer A + 0.01% Tween, 2X in Dynabeads buffer B + 0.01% Tween, 3X in Ribohyb buffer and resuspended in a final volume of 10 ul Ribohyb buffer in a non-stick tube (Eppendorf Protein LoBind). The 10 µl hybridization reactions were added to 10 µl of resuspended beads for a binding reaction volume of 20 µl. Samples were then incubated in a thermomixer at 50 °C for 10 min at 1000 rpm, then moved to a magnetic rack. The supernatant (rRNA depleted) was taken, the beads were washed with 20 µl of Ribowash buffer (0.4X SSC, 0.01% Tween), and the second supernatant combined with the first supernatant. The supernatant was diluted 1:8 in water and 4 µl of the dilution was used for cDNA synthesis.

### rRNA removal with pooled probes

A mix of 15 oligos targeting the 18S and 28S were biotinylated. For low-input reactions, 4 pmol probe mix was combined with 0.5 or 100 ng of total RNA along with 4U µl RNasin Plus (Promega) and Ribohyb buffer. Reactions were denatured at 70°C for 5 min and then added directly to 4 µl of prewashed MyOne C1 streptavidin beads with the supernatant removed such that the volume of the binding reaction remained the same as the hybridization reaction. Beads were prepared as for the single-probe assay, except Tween was withheld from Dynabeads buffer A. (We observed that Tween forms precipitates in buffer A after storage for > 1 week and thus decided to omit it for further experiments.) After binding, the supernatant was collected and the beads were washed with 10 µl of Ribowash buffer and the supernatant collected again. The 20 µl of Ribo-Pop depleted supernatant was cleaned up with 1.8X volume of Mag-Bind TotalPure NGS (Omega Bio-tek) beads to recover total RNA. High-input reactions containing 1 µg of total RNA were mixed with 40 pmol probe mix and 40U RNasin Plus in a 50 µl reaction. The reactions were mixed with 40 ul of beads and performed otherwise identically to the low-input reactions but with 5X higher volume for washing and RNA cleanup. For the samples used for RNA-seq, 5 µg of total RNA was mixed with 200 pmol probe mix and 40U RNasin Plus in a 40 µl reaction. The binding reaction was done in a total volume of 50 µl in Ribohyb buffer and after binding, beads were washed once with 50 µl Ribowash buffer and the supernatants with cleaned up with Mag-Bind beads. After cleanup, Ribo-Pop depleted RNA was subjected to cDNA synthesis and qPCR analysis, where it was compared to a non-depleted input RNA sample from the same source material.

### RNA sequencing and data analysis

Ribo-Pop rRNA removal was applied to 5 µg of larval RNA as described in the previous section. A synthetic RNA spike-in mix, Lexogen E0 (Lexogen, cat # 025.03) was added to the sample before rRNA removal at a ratio of 0.09 ng per 100 ng total RNA for quality control. After rRNA removal, fractions of both the input RNA and the rRNA-depleted RNA were used to construct total RNA libraries or poly(A)-primed libraries. We used QuantSeq 3′ mRNA-Seq Library Prep Kit FWD for Illumina (Lexogen, cat # 015.24) to prepare poly(A)-primed libraries. For the input samples, 500 ng of total RNA was used for library construction. For rRNA-depleted samples, the equivalent amount RNA corresponding to 500 ng of total RNA pre-depletion was used (11 ng). Libraries were amplified with 12 cycles of PCR. We used the NEBNext Ultra II Directional RNA Library Prep Kit (E7765S) to construct the total RNA libraries. We used the equivalent amount of RNA corresponding to 1 µg of total RNA pre-depletion (22 ng) or a matched amount of non-depleted total RNA. Libraries were amplified for 8 PCR cycles. Libraries were loaded onto an Illumina NextSeq 500/550 High Output v2 cartridge and subjected to single-end sequencing on a NextSeq 500 instrument for 85 cycles.

The raw sequencing data was de-multiplexed and converted to fastq format on Illumina Basespace. First adapters were removed from the sequences. For the NEB libraries, the NEB adapter was removed with Cutadapt^40^. For the QuantSeq libraries, poly(A) and Illumina adapters were trimmed from the reads using BBDuk^41^ with the parameters “k=13 ktrim=r useshortkmers=t mink=5 qtrim=r trimq=10 minlength=20”, as recommended by Lexogen. *Drosophila melanogaster* genomic and transcript sequences were downloaded from Ensembl (release 99, assembly BDGP6.28). A STAR^42^ index was built containing the genomic sequence and the SIRV synthetic spike-in gene sequences (Lexogen). A Kallisto^43^ index was built with the coding, non-coding, and spike-in transcripts, as well intronic sequences with 30 nt of flanking sequence on either side. Kallisto was used to estimate transcript abundance from all libraries using the “-l 1 -s 1 --single --fr-stranded” options for the QuantSeq libraries and “-l 1 -s 1 --single --rf-stranded” for the NEB libraries. Gene-level differential expression analysis was performed with DESeq2 using the transcript import function^44^. Reads were also mapped to the genome with STAR to obtain alignments for visualizing coverage on genomic regions. The coverage of the rRNA locus presented in Fig. 5 was obtained using Samtools^45^ and Bedtools^46^. After sorting the alignments with Samtools, Bedtools was used to calculate coverage using the -bga option to produce a bedgraph file. Gene-level quantification presented in Fig. 4 was obtained by summing quantification of individual transcripts. For biotype analysis, most genes were assigned to the biotype given in the annotation file. An exception was made for pseudogenes derived from rRNA, which were assigned to the rRNA biotype. Snakemake^22^ and Conda^47^ were used to run the RNA-seq pipeline and manage software dependencies. Downstream analysis and figure construction was completed using open-source scientific computing software, including Numpy, Scipy, Pandas, and Matplotlib, run in Jupyter notebooks^48–52^. Illustrations were made in Inkscape. Further details, including the RNA-seq pipeline and details of downstream analysis are available on Github at https://github.com/marykthompson/ribopop_rnaseq. The specific versions of software used are listed in Table S3.

## Supporting information

Supplemental protocol

## DATA DEPOSITION

RNA sequencing data, including processed files summarizing the gene-level quantification and differential expression analysis, have been deposited in the Gene Expression Omnibus with accession number GSE150332.

## SUPPLEMENTAL MATERIAL

Supplemental figures 1 – 4

Supplemental protocol

Table S1: Sequences and characteristics of the 35 probes individually tested for 18S depletion

Table S2: Pre-designed Ribo-Pop probe sets for *Drosophila*, yeast (*S. cerevisiae* & *S. pombe*), and mammal (*H. sapiens* & *M. musculus*)

Table S3: Additional oligo sequences and software versions

## ACKNOWLEDGMENTS

This work was funded by a Wellcome Investigator Award 209412/Z/17/Z and a Leverhulme Trust Research Project Grant to I.D. M.K.T. was also funded by the European Union’s Horizon 2020 research and innovation programme under Marie Skłodowska-Curie grant agreement 750928. M.K. was supported by the Biotechnology and Biosciences Research Council (BBSRC), grant numbers: (BB/M011224/1) and (BB/S507623/1) and by Zegami Ltd.

## Author contributions

M.K.T. designed the study, performed experiments, analyzed the data, and wrote the manuscript. I.D. advised on experiments and wrote the manuscript. M.K. contributed to the computational probe design pipeline.

---

**FIGURE S1.**
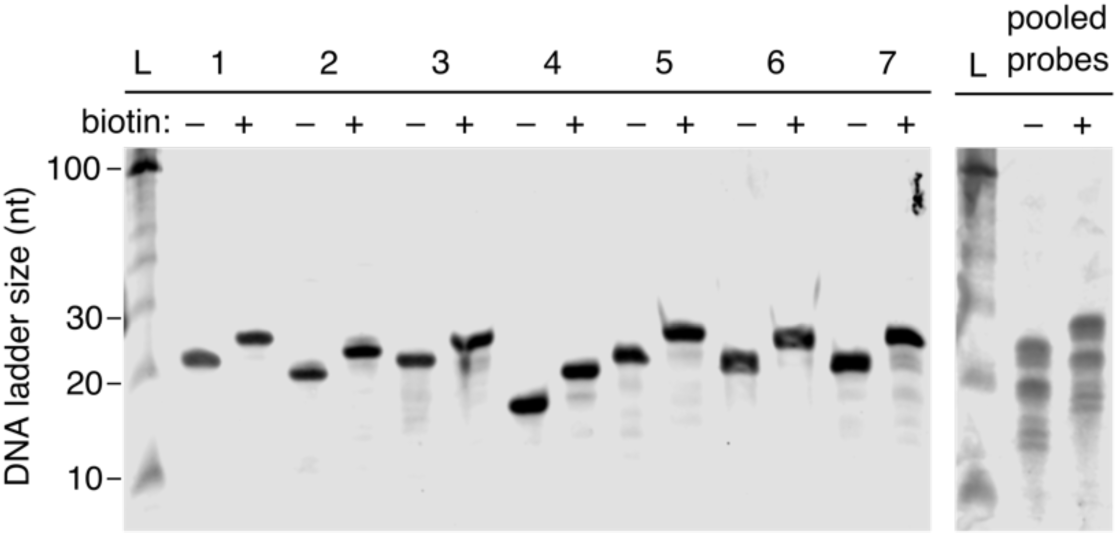
Enzymatic addition of a single 3′ biotin moiety to unmodified oligos. Unmodified oligos were biotinylated with biotin-ddUTP and terminal deoxynucleotidyl transferase (TdT). Biotinylation efficiency was assessed by Urea PAGE. Successful biotinylation causes an upward shift of the oligo in the gel. In each lane, 1 pmol of oligo was loaded. Left: Biotinylation efficiency assessment for single-probe reactions, a representative gel is shown. Right: Biotinylation efficiency assessment for the pool of 15 oligos. L = DNA ladder.

**FIGURE S2.**
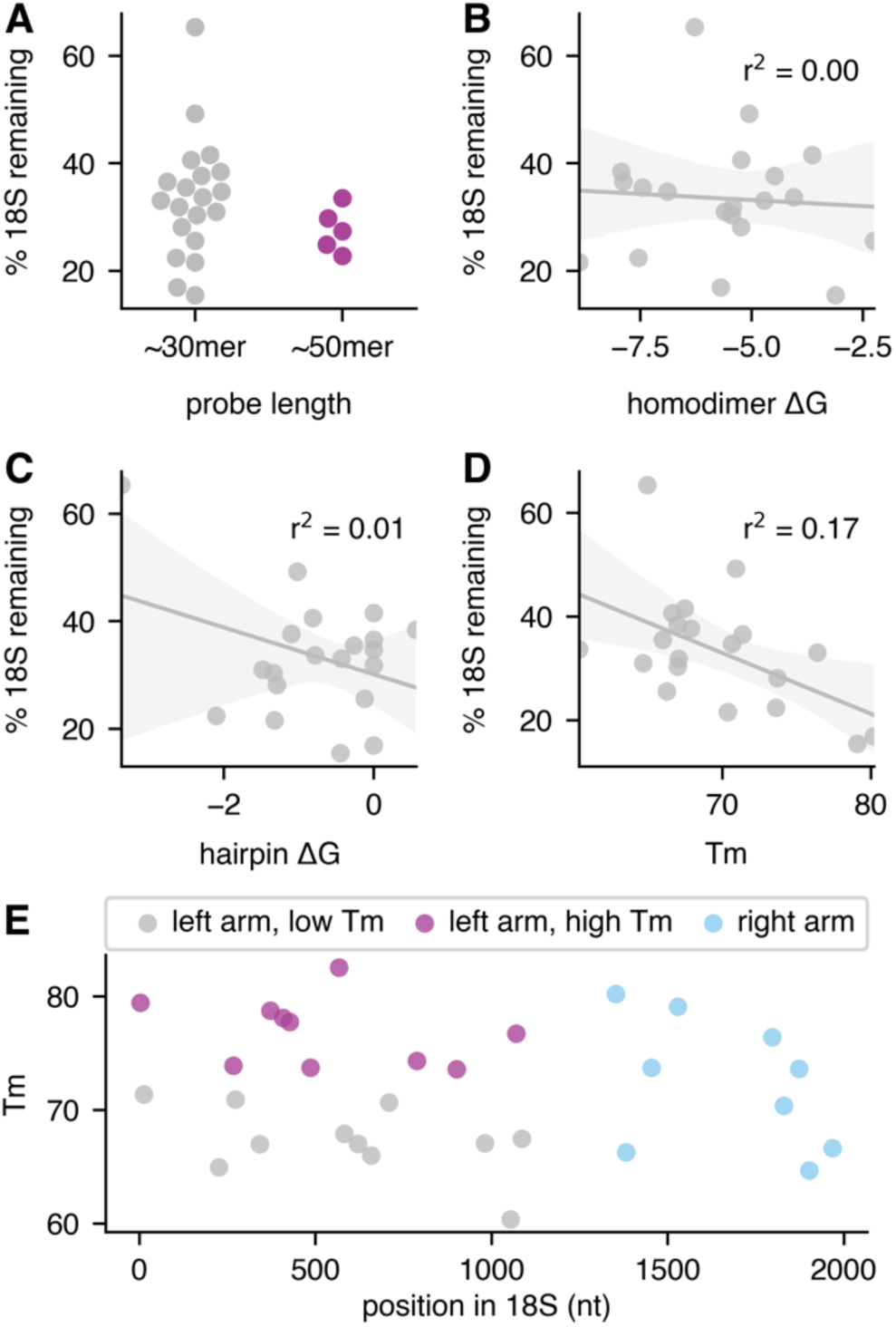
Correlation of probe properties with performance in the single-probe depletion assay. A) Comparison of depletion values for short probes (26 – 32 nt) and longer probes (46 – 52 nt). Two-sided t-test p = 0.38. B-D) Depletion values vs. predicted thermodynamic properties of the antisense oligos, including probe self-structure (B & C) and probe/target melting temperature (D). P-values for the Spearman’s correlations are 0.99, 0.65, and 0.07 for B-D. E) Predicted Tm of the tested probes plotted by target position in the 18S rRNA.

**FIGURE S3.**
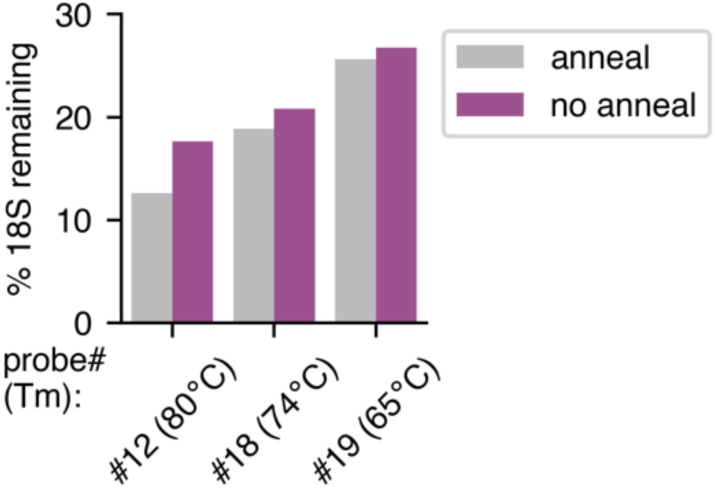
Removal of the annealing step does not affect rRNA depletion. Three different probes were tested for their ability to deplete 18S rRNA from total RNA with or without a 10 minute annealing step after denaturation (allowing the reaction to cool slowly to room temperature before binding to the streptavidin beads). The values shown are the averages of technical qPCR replicates.

**FIGURE S4.**
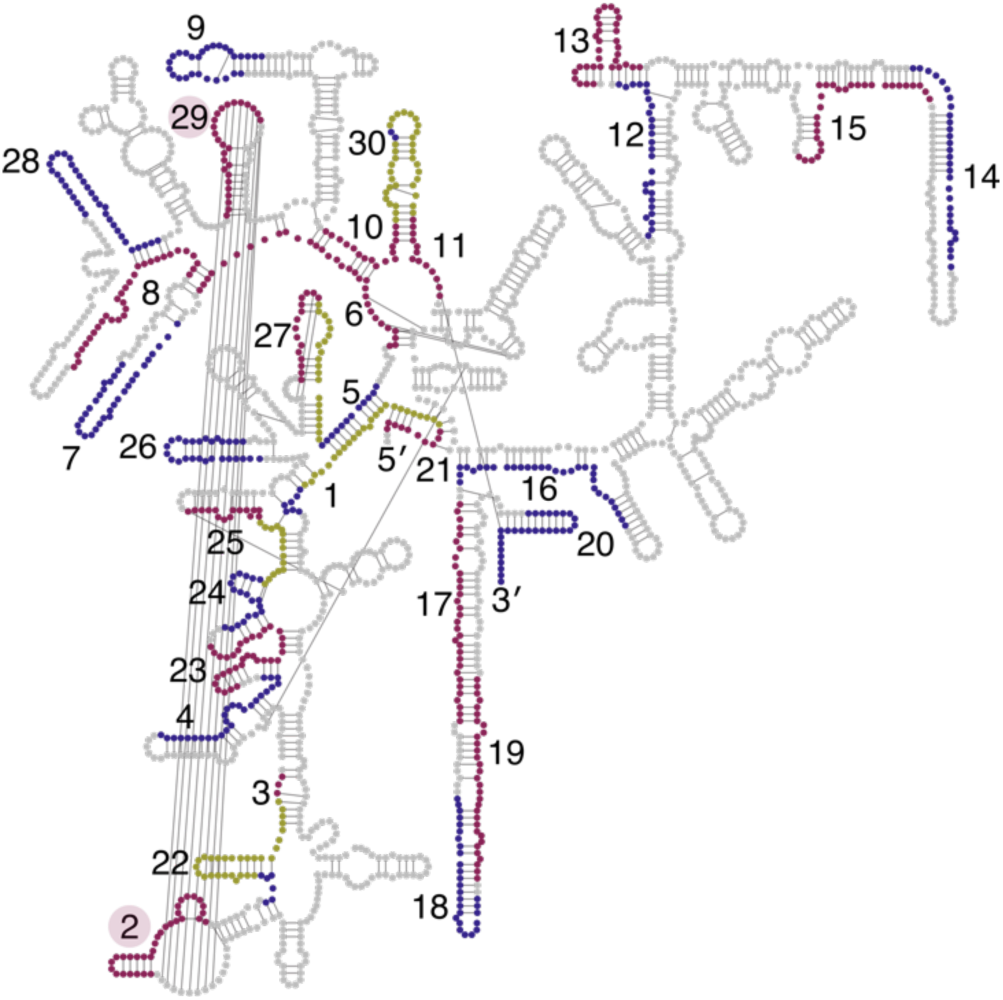
Probe targeting in heavily structured 18S regions. Target sites of tested probes in the *Drosophila* 18S rRNA, as in Fig. 2, but with the detected base-pairing interactions shown. Note the interaction between the target sites of probe #2 and probe #29. Some obvious base-pairing in helices is absent from the annotated base pairs due to the method of assigning pairing, which was derived from the 3D structure file^13^.

