## Supplemental protocol for "Ribo-Pop: Simple, cost-effective, and widely applicable ribosomal RNA depletion"

**Supplemental protocol: rRNA depletion from high or low input samples**

**Prepare MyOne C1 beads:**

1. Mix the bead stock and aliquot the calculated volume to a non-stick tube.
2. Batch wash the beads in 2X original volume, or at least 100 µl.
   i. Wash 3X in 1X MyOne B&W buffer + Tween.
   ii. Wash 2X in Dynabeads buffer A.
   iii. Wash 2X in Dynabeads buffer B + Tween.
   iv. Wash 2X in 1X Ribohyb buffer.
3. Resuspend beads in 5X original volume of 1X Ribohyb buffer and aliquot 5X per-sample bead volume to a non-stick tube for each sample. Leave the prepared beads in a magnetic rack until ready to proceed.

**Probe hybridization and streptavidin pulldown*:**

1. Add the following components to a tube on ice (an example is shown for 5 µg RNA, scale down to as low as 10 µl for low input reactions):

| component | µl |
| --- | --- |
| probe mix, 20 pmol/µl | 10 |
| total RNA, 200 ng/µl | 25 |
| 10X Ribohyb buffer | 4 |
| RNAse inhbitor (optional) | 1 |
| H2O to 40 µl | 0 |

2. Move the hybridization reactions to a heat block or thermomixer at 70 ˚C for 5 min.
3. Remove the supernatant from the prepared MyOne C1 beads and add the hybridization reaction. Mix thoroughly by pipetting, making sure to fully resuspend the beads.
4. Incubate the reaction at 50 ˚C for 10 min with vigorous mixing, for example using a Thermomixer at 1000 rpm.
5. Place the tubes in the magnetic stand and remove the supernatant to a new tube (save the supernatant!).
6. Add 1 reaction volume of Ribowash buffer, resuspend the beads, place back on the magnet, and take the supernatant again. Add the 2^nd^ supernatant to the first supernatant.
7. Proceed to clean up of the combined supernatants.

*Notes:

- Calculation of probe molar excess: Using the length of the major rRNA transcripts and the average MW of 340 gm/mol*nt, we estimate that 200 ng of total RNA contains 0.2 pmol of rRNA (approximately 0.1 pmol of 18S and 0.1 pmol of 28S). We need 0.5 pmol of each probe to reach a 5-fold molar excess. In a 15-probe pool, this is 7.5 pmol of pooled probe mix. Scaled to 5 µg total RNA, this becomes 187.5 pmol, which we rounded to 200 pmol for simplicity.
- Calculation of bead excess: We achieved good results using a 5:1 ratio of stated bead binding capacity to oligo. For the MyOne C1 beads, the stated binding capacity for oligos is 500 pmol/mg. At 10 mg/ml, the beads can bind 5 pmol/µl. Therefore, we simply used 200 µl of beads (1000 pmol binding capacity) to capture 200 pmol of probe.
- It is difficult to know without testing what ratio of probe:RNA should be used for a new sample type or probe set. We have observed that both decreasing the reaction volume (and therefore increasing the probe and target concentrations) and increasing the molar excess of probe increases the efficiency of rRNA removal. We recommend choosing the smallest hybridization reaction possible given the starting concentration of the total RNA (down to 10 µl). Then, test reactions can be performed starting with 5-fold molar excess of probe:RNA and compared to an aliquot of the input RNA by qPCR. Based on our sequencing results, we believe that for most successful depletion it is desirable to see target depletion at <1% of input levels by qPCR.
- These reactions should not be transferred to ice at any of these steps, in case cooling could cause non-specific binding of the probes.
- It may be helpful to briefly spin the reactions after the denaturation step and the binding step. The spin should be done in a non-refrigerated centrifuge, for example a benchtop microcentrifuge.

**Clean up the supernatant:**

RNA can be purified from the supernatant with alcohol precipitation, a column-based clean-up kit, or a bead-based clean-up kit. We recommend using a column- or bead-based kit, as the standard protocol should also remove any remaining small DNA oligos. Remaining oligo can be assessed by running an aliquot of the rRNA depleted and purified reaction alongside an aliquot of the oligo mix on a PAGE gel (e.g. 20% TBE-Urea). We were not able to observe any remaining oligo by PAGE after streptavidin pulldown and bead-based clean up.

**TdT labeling of unmodified oligos (required if starting with unmodified oligos)^1^:**

1. Dilute individual unmodified oligos to 200 µM in probe resuspension buffer.
2. Add an equal volume of each unmodified oligo to obtain a 200 µM pool.
3. Set up the TdT labeling reaction in a PCR tube with the following components:

| component | µl |
| --- | --- |
| oligo mix, 200 µM | 5 |
| biotin-ddUTP, 1 mM | 3 |
| 5X TdT buffer | 3 |
| H2O | 3.4 |
| TdT enzyme (20 U/µl) | 0.6 |

4. Incubate overnight at 37 ˚C in a PCR machine.
5. Stop the reaction by heating to 70 ˚C for 10 min.
6. Clean up the oligos using ethanol precipitation or an oligo clean-up kit (e.g. Zymo Oligo Clean & Concentrator).

**Buffers:**

10X Ribohyb buffer*:

20X SSC, 0.1% Tween-20

1X Ribowash buffer:

0.4X SSC, 0.01% Tween-20

20X SSC:

3 M NaCl, 300 mM trisodium citrate (adjust pH to 7.0 with HCl).

Probe resuspension buffer:

10 mM Tris-HCl (pH 8.0), 0.1 mM EDTA

*Use 10X for setting up the hybridization reactions and make a 1X dilution for bead preparation.

Dynabeads buffers (from MyOne C1 protocol with added Tween-20):

1X MyOne B&W buffer + Tween:

5 mM Tris-HCl (pH 7.5), 0.5 mM EDTA, 1 M NaCl, 0.01% Tween-20

Dynabeads buffer A:

0.1 M NaOH, 0.05 M NaCl

Dynabeads buffer B + Tween:

0.1 M NaCl + 0.01% Tween-20
